# Microbial phylogenetic relatedness links to distinct successional patterns of bacterial and fungal communities

**DOI:** 10.1101/2021.05.19.444715

**Authors:** Qiang Lin, Francisco Dini-Andreote, Travis B. Meador, Roey Angel, Lenka Meszárošová, Petr Heděnec, Lingjuan Li, Petr Baldrian, Jan Frouz

**Affiliations:** Biology Centre of the Czech Academy of Sciences, Institute of Soil Biology & SoWa Research Infrastructure, Na Sádkách 7, CZ, 37005, České Budějovice, Czech Republic; Department of Plant Science, The Pennsylvania State University, University Park, PA, USA; Huck Institutes of the Life Sciences, The Pennsylvania State University, University Park, PA, USA; Laboratory of Environmental Microbiology, Institute of Microbiology of the Czech Academy of Sciences, Vídeňská 1083, 14220 Praha 4, Czech Republic; Institute for environmental studies, Faculty of Science, Charles University, Benátská 2, 12800, Praha 2, Czech Republic; Department of Geosciences and Natural Resource Management, Faculty of Science, University of Copenhagen, Rolighedsvej 23, 1958 Frederiksberg C, Denmark; Engineering Research Center of Soil Remediation of Fujian Province University, College of Resources and Environment, Fujian Agriculture and Forestry University, Fuzhou 350002, China

**Author notes:** **Corresponding author**: Qiang Lin, and Jan Frouz.

**Keywords:** soil bacteria and fungi, ecological succession, community assembly, phylogenetic relatedness, co-occurrence, post-mining lands

## Abstract

Development of soil microbial communities along ecological succession is crucial for ecosystem recovery and maintenance. However, mechanisms mediating microbial community dynamics and co-occurrence patterns along ecological succession remain unclear. Here, we explored community dynamics and taxa co-occurrence patterns in bacterial and fungal communities across a well-established chronosequence of post-mining lands spanning 54 years of recovery. Meanwhile, by synthesizing previous studies and ecological theories, we devised two conceptual models that integrate microbial phylogeny with patterns in community dynamics and in taxa co-occurrence. We further tested these models by using empirical data. At early successional stages, bacterial community structures became increasingly phylogenetically clustered with soil age, which was co-determined by the environmental selection from soil vegetation cover and by heterogeneous responses of less phylogenetically similar bacteria to the increasing resource availability along succession. At later successional stages, bacterial community phylogenetic structures displayed progressively lower variability. The fungal community phylogenetic structures varied relatively less and were independent of soil age, soil properties and vegetation cover, which was attributed to the dominance of stochastic processes in community turnover along succession. Network analysis revealed a decrease in bacterial co-occurrence complexity along succession, which aligned with a decrease in average pairwise phylogenetic distances between co-occurring bacteria. These patterns together implied a decrease in potential bacterial cooperation that was probably mediated by increasing resource availability along succession. The increased complexity of fungal co-occurrence along succession was independent of phylogenetic distances between co-occurring fungi. This study provides new sights into ecological mechanisms underlying bacterial and fungal community succession.

## 1. Introduction

Elucidating the co-development of soil microbiomes and the ecosystem that hosts them is crucial for understanding ecosystem functioning, recovery and maintenance (Bardgett and van der Putten, 2014; Dini-Andreote et al., 2014; Bahram et al., 2018). Therefore, studying natural development of post-mining lands provides a unique opportunity to advance knowledge on the mechanisms underlying soil microbiome development along with *de novo* ecosystem recovery, as strip-mining activities often degrade existing ecosystems to an almost primordial stage.

Elucidating soil microbial community dynamics and microbial taxa co-occurrence patterns along ecological succession could provide important clues to understand successional patterns of microbial communities (Brown and Jumpponen, 2014; Dini-Andreote et al., 2014; Harantová et al., 2017; Morriën et al., 2017), whereas addressing either of them remains challenging (Additional file 1: Fig. S1). Besides environmental selection effects, the niche characteristics of the initial community could also affect community dynamics (i.e., the transition from an initial community to a later community), as community niche characteristics determine how the community respond to environmental selection/disturbance. For example, a few studies have revealed “legacy effects” based on the finding that initial microbial communities differing in niche characteristics, subsequently subjected to the identical environmental selection, finally transitioned to different community assemblages (Evans and Wallenstein, 2014; Banerjee et al., 2016a). However, it remains challenging to understand how the niche characteristics of the initial community would affect community dynamics (Additional file 1: Fig. S1a).

Due to the high complexity of soil microbiomes, direct evaluation of their niche characteristics is not technically feasible. Instead, community phylogenetic structures could provide clues for community niche characteristics (Webb et al., 2002). As such, a phylogenetically clustered community (species in this community are more closely related than null model expectation (Stegen et al., 2012)) consists of species with similar niches; an overdispersed community (species in this community are more distantly related than null model expectation (Stegen et al., 2012)) consists of species with distinct niches (Webb et al., 2002). Here, to address the challenge in elucidating community dynamics (Additional file 1: Fig. S1a), we devised a conceptual model (Fig. 1a) to describe how the phylogenetic structure of the initial community will affect the later community structure, namely community dynamics. In our model (Fig. 1a), if the initial community is phylogenetically overdispersed (nearest taxon index (NTI) < −2 (Stegen et al., 2012)), this community is predicted to show heterogeneous responses to the environmental selection. In this case, the later community is expected to become more phylogenetically clustered than the initial community. For example, Tripathi et al. found that under pH selection, the phylogenetically overdispersed (or less phylogenetically clustered) community became more phylogenetically clustered (Tripathi et al., 2018). This happens because taxa performed heterogeneous responses that well-adapted taxa became predominant while maladapted taxa became rare or were excluded in the later community. These predominant taxa are probably closely related, as they show similar adaptations to pH selection (Webb et al., 2002), thus resulting in a more phylogenetic clustered community. In contrast, if the initial community is phylogenetically clustered (nearest taxon index (NTI) > 2 (Stegen et al., 2012)), this community is predicted to show a homogeneous response to the environmental selection (Fig. 1a), so that the phylogenetic structure of the later community is expected to keep relatively stable as the initial community. This happens because taxa perform heterogeneous responses that taxa in the initial community are closely related and thus are presumed to develop similarly in response to the environmental selection (Webb et al., 2002). Taken together, our model (Fig. 1a) conceptualizes another factor (i.e., the phylogenetic structure of the initial community) underlying community dynamics along ecological succession.

**Fig. 1.**
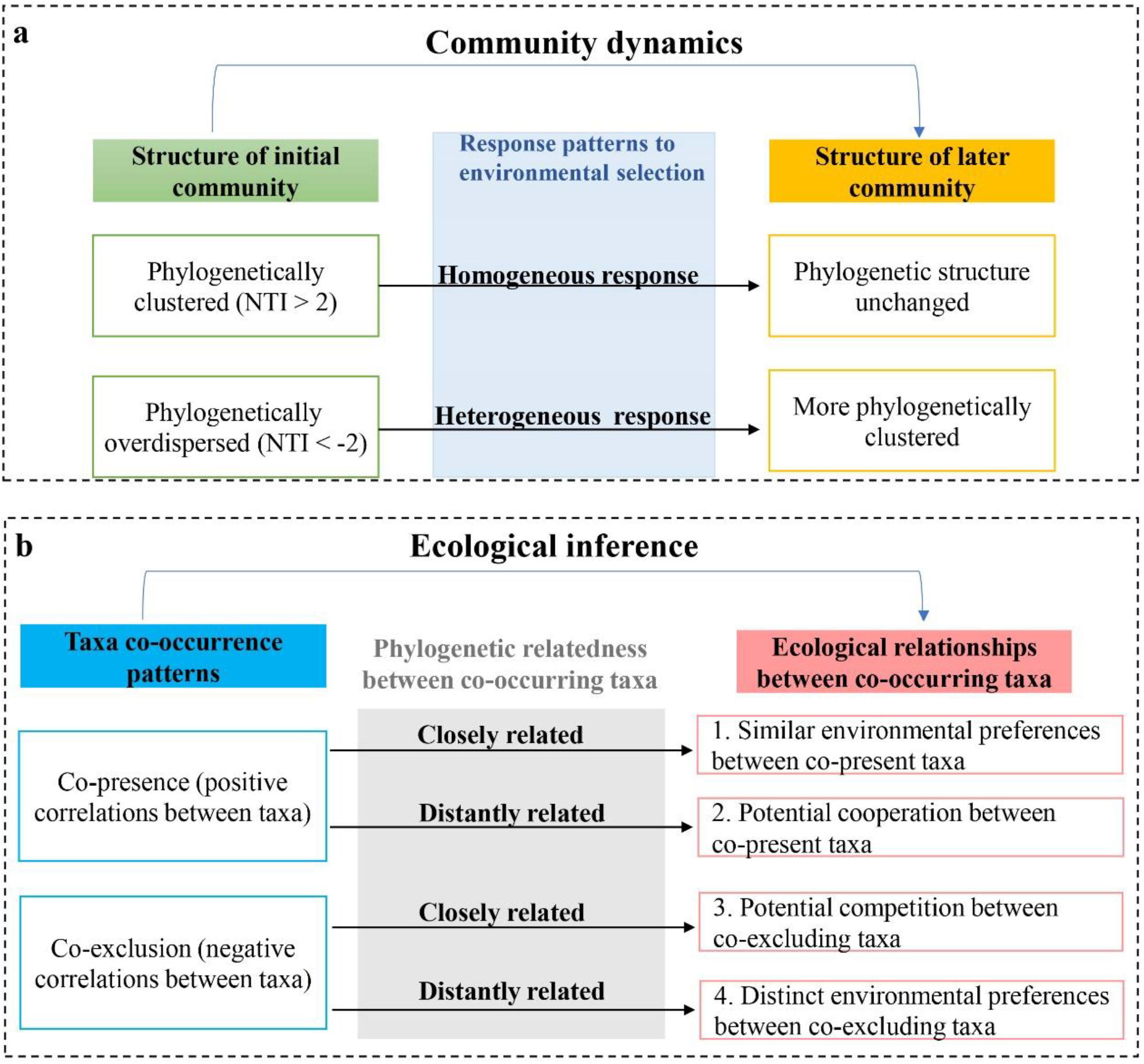
Conceptual models in elucidating microbial community dynamics (a) and taxa co-occurrence (b) separately. The model (a) uses the phylogenetic structure of the initial community to predict the structure of the later community, namely community dynamics. If the initial community is phylogenetically overdispersed (nearest taxon index (NTI) < −2), this initial community is predicted to show heterogeneous responses to the environmental selection, so that the later community is expected to become more phylogenetically clustered than the initial community. If the initial community is phylogenetically clustered (NTI > 2), this initial community is predicted to show a homogeneous response to the environmental selection, so that the phylogenetic structure of the later community is expected to keep relatively stable as the initial community. The model (b) integrates taxa co-occurrence patterns and phylogenetic relatedness between the co-occurring taxa to differentiate ecological inferences from taxa co-occurrence. Taxa co-presence and co-exclusion are separately based on positive and negative correlations between taxa in a network. Taxa phylogenetic relatedness is evaluated by phylogenetic distance between co-occurring taxa.

Although microbial taxa co-occurrence patterns are important for understanding microbial community assembly along ecological succession (Dini-Andreote et al., 2014; Morriën et al., 2017), ecologically explaining taxa co-occurrence patterns is challenging (Additional file 1: Fig. S1b). For example, taxa co-presence (positive correlations between taxa in the network) could imply two distinct ecological inferences: 1) potential cooperation for survival/development across taxa with distinct niches, or 2) similar environmental preferences across taxa with similar niches (Barberan et al., 2012; Deng et al., 2012). Taxa co-exclusion (negative correlations between taxa) could also imply two distinct ecological inferences: 1) potential competition across taxa with similar/overlapped niches, or 2) distinct environmental preferences across taxa with distinct niches (Barberan et al., 2012; Deng et al., 2012) (Additional file 1: Fig. S1b). Therefore, it is challenging to differentiate the ecological inferences derived from taxa co-occurrence patterns and thus hinders the understanding of ecological relationships among taxa (Additional file 1: Fig. S1b).

Remarkably, all ecological inferences derived from microbial co-occurrence patterns are tightly linked to species niches, so the phylogenetic relatedness among co-occurring species can help to differentiate the ecological inferences derived from a specific co-occurrence pattern, based on the assumption of phylogenetic niche conservatism (Webb et al., 2002). Here, to address the challenge in ecologically elucidating taxa co-occurrence patterns (Additional file 1: Fig. S1b), we devised a conceptual model (Fig. 1b) that integrates phylogenetic relatedness between the co-occurring taxa with taxa co-occurrence patterns to differentiate ecological inferences from microbial co-occurrence. In the model (Fig. 1b), in scenario (1): taxa co-presence (positive correlations between taxa in the network) implies similar environmental preferences between the co-present taxa, if the co-present taxa are closely related (evaluated by the phylogenetic distance between the co-present taxa), because similar environmental preferences are usually found among closely related taxa (Webb et al., 2002); in scenario (2): taxa co-presence implies potential cooperation between the co-present taxa, if the co-present taxa are distantly related, because functionally complementary cooperation tends to establish between distantly related taxa (Morris et al., 2013; Zelezniak et al., 2015); in scenario (3): taxa co-exclusion (negative correlations between taxa in the network) implies potential competition between the co-excluding taxa, if the co-excluding taxa are closely related, because competition tends to occur between closely related taxa that occupy similar/overlapped niches (Hibbing et al., 2010); in scenario (4): taxa co-exclusion implies distinct environmental preferences between the co-excluding taxa, if the co-excluding taxa are distantly related, because distinct environmental preferences are usually found among distantly related taxa (Webb et al., 2002).

In this study, we focused on two major components of soil microbiomes (bacterial and fungal communities) and utilized phylogenetic information to disentangle the interplay of ecological processes underlying microbial community dynamics and co-occurrence patterns along ecological succession on post-mining lands. We addressed three main questions: (*i*) To what extent do the phylogenetic structures of bacterial and fungal communities change along succession? (*ii*) What are the relative influences of distinct ecological processes that mediate bacterial and fungal community dynamics? (*iii*) How do bacterial and fungal co-occurrence patterns change along ecological succession, and why?

## 2. Materials and methods

### 2.1. Data collection

A well-established soil chronosequence from the lignite mining district near Sokolov in the Czech Republic (Frouz et al., 2001; Frouz and Nováková, 2005; Mudrak et al., 2016) was used to investigate the patterns of soil succession and recovery. The chronosequence and other background information in this area were provided by Sokolovská Uhelná mining company. The chronosequence was further validated using historical aerial photography and additional independent methods (Frouz, 2013). In this area, the mean annual temperature is 6.8 °C, the mean annual precipitation is 650 mm and the altitude is ∼550 m a.s.l (Frouz et al., 2001). This chronosequence spans 54 years of ecosystem development from bare overburden of alkaline Miocene clay via herbs, grasses, goat willow shrubs to young forests dominated by birch and aspen (Mudrak et al., 2016; Harantová et al., 2017). A total of four distinct chronosequence sites (i.e., Sites I, II, III and IV) were sampled in triplicate at three time-points (in the end of May in 2006 (2008 only for the Site I), 2010 and 2015) (Additional file 1: Table S1) (see (Harantová et al., 2017) for additional details). The aboveground vegetation was surveyed as previously described (Harantová et al., 2017). Soil samples were subjected to total DNA extraction (Additional file 2: S1), quantification of microbial biomass, and characterization of physicochemical properties (i.e., total nitrogen, organic carbon and pH), as previously described (Sagova-Mareckova et al., 2008; Harantová et al., 2017). The primer sets 515F/806R (Caporaso et al., 2012) and nu-SSU-0817/nu-SSU-1196 (Borneman and Hartin, 2000) were used to amplify the bacterial 16S rRNA gene and the fungal 18S rRNA gene, respectively, as previously described (Zifcakova et al., 2016; Navrátilová et al., 2019). PCR products were purified and sequenced on an Illumina MiSeq platform in the Laboratory of Environmental Microbiology, Institute of Microbiology of the Czech Academy of Sciences.

### 2.2. Data analysis

Amplicon sequences were sorted according to their unique barcodes and trimmed for quality (sequences containing ambiguous characters, with read length < 200bp, or with quality score < 15 were removed) (Edgar and Flyvbjerg, 2015), followed by chimera removal (Edgar et al., 2011). Sequences were clustered at 97% and 98.5% similarities, into operational taxonomic units (OTUs), for bacteria and fungi, respectively (Li and Godzik, 2006). Taxonomic assignments of bacterial sequences were performed using the Greengenes database (version 13_8) (DeSantis et al., 2006). Each fungal OTU was assigned to its closest genus using the Genbank database. Singletons, non-bacterial and non-fungal sequences were removed, and the OTU tables were rarefied to 7942 sequences per sample (bacteria) and 1431 sequences per sample (fungi).

Representative sequences of fungal and bacterial OTUs were aligned with PyNAST (Caporaso et al., 2010), referring to the SILVA alignment version 108 (https://www.arb-silva.de/download/archive/qiime/), and the Greengenes core set alignment (http://greengenes.lbl.gov/Download/Sequence_Data/Fasta_data_files/). The high-quality alignments of bacterial and fungal representative sequences were used to construct phylogenetic trees with FastTree, respectively (Price et al., 2009). To evaluate the phylogenetic relatedness between OTUs, the phylogenetic distance of pairwise OTUs was determined using the function *cophenetic* in the R (v. 4.0.2) package ‘stats’ (v. 4.0.2) based on the phylogenetic tree. To evaluate the niche differences between OTUs, the niche distance of pairwise OTUs for all measured environmental variables (aboveground vegetation and soil properties) was determined as previously described (Stegen et al., 2012). To test the phylogenetic signal, the function *mantel*.*correlog* (permutations = 999) in the R package ‘vegan’ (v. 2.5-6) was used to measure the Pearson’s correlation between OTU phylogenetic distances and OTU niche distances (Stegen et al., 2012). To evaluate the community phylogenetic structure, nearest taxon index (NTI) of microbial communities was calculated in the R package ‘picante’ (v.1.8.2) with the function *ses*.*mntd* (abundance.weighted = TRUE, null.model=“taxa.labels”, iterations = 1000) with 999 randomization across all samples. NTI value > +2 indicates phylogenetic clustering (species in a local community are more closely related than null model expectation), whereas NTI value < −2 indicates phylogenetic overdispersion (species in a local community are more distantly related than null model expectation) (Table 1) (Stegen et al., 2012; Stegen et al., 2013). Mantel test was used to evaluate correlations between Euclidean distances of NTI values and vegetation cover, using the R package ‘vegan’. To estimate pairwise phylogenetic turnover between communities, Beta mean nearest taxon distance (βMNTD) was calculated using the function *comdistnt* (abundance.weighted = TRUE) in the R package ‘picante’. β-nearest taxon index (βNTI) was used to estimate the degree to which observed βMNTD deviates from the mean of the null distribution βMNTD, normalized by its standard deviation. βNTI was calculated based the phylogenetic null model (Additional file 2: S2) (Stegen et al., 2013; Stegen et al., 2015). βNTI value < −2 indicates that phylogenetic turnover is driven by homogeneous selection (significantly less turnover than null model expectation), whereas βNTI value > +2 indicates variable selection (significantly greater turnover than null model expectation) (Table 1) (Stegen et al., 2013; Stegen et al., 2015). |βNTI| < 2 indicates that phylogenetic turnover is not driven by deterministic processes (no significant differences between observed turnover and null model expectation) but by dispersal limitation, homogenizing dispersal, or undominated process (including ecological drift and other stochastic processes excluding dispersal) (Table 1) (Stegen et al., 2013; Stegen et al., 2015). In the scenario of |βNTI| < 2, to disentangle these ecological processes, the Bray-Curtis-based Raup-Crick metric (RC_bray_) was calculated with the modified approach by Stegen et al. (Stegen et al., 2013) from Chase et al. (Chase et al., 2011). Taken together, the relative importance of ecological processes to the phylogenetic turnover of microbial communities was evaluated following the previous method (Table 1) (Stegen et al., 2013; Stegen et al., 2015). In particular, in the scenario of |βNTI| < 2, the percentages of RC_bray_ > +0.95, RC_bray_ <− 0.95 and |RC_bray_| < 0.95 were used to quantify the relative importance of dispersal limitation, homogenizing dispersal and undominated process, respectively. The percentages of βNTI values > +2 and < −2 were used to quantify the relative importance of variable and homogeneous selection, respectively.

**Table 1.**
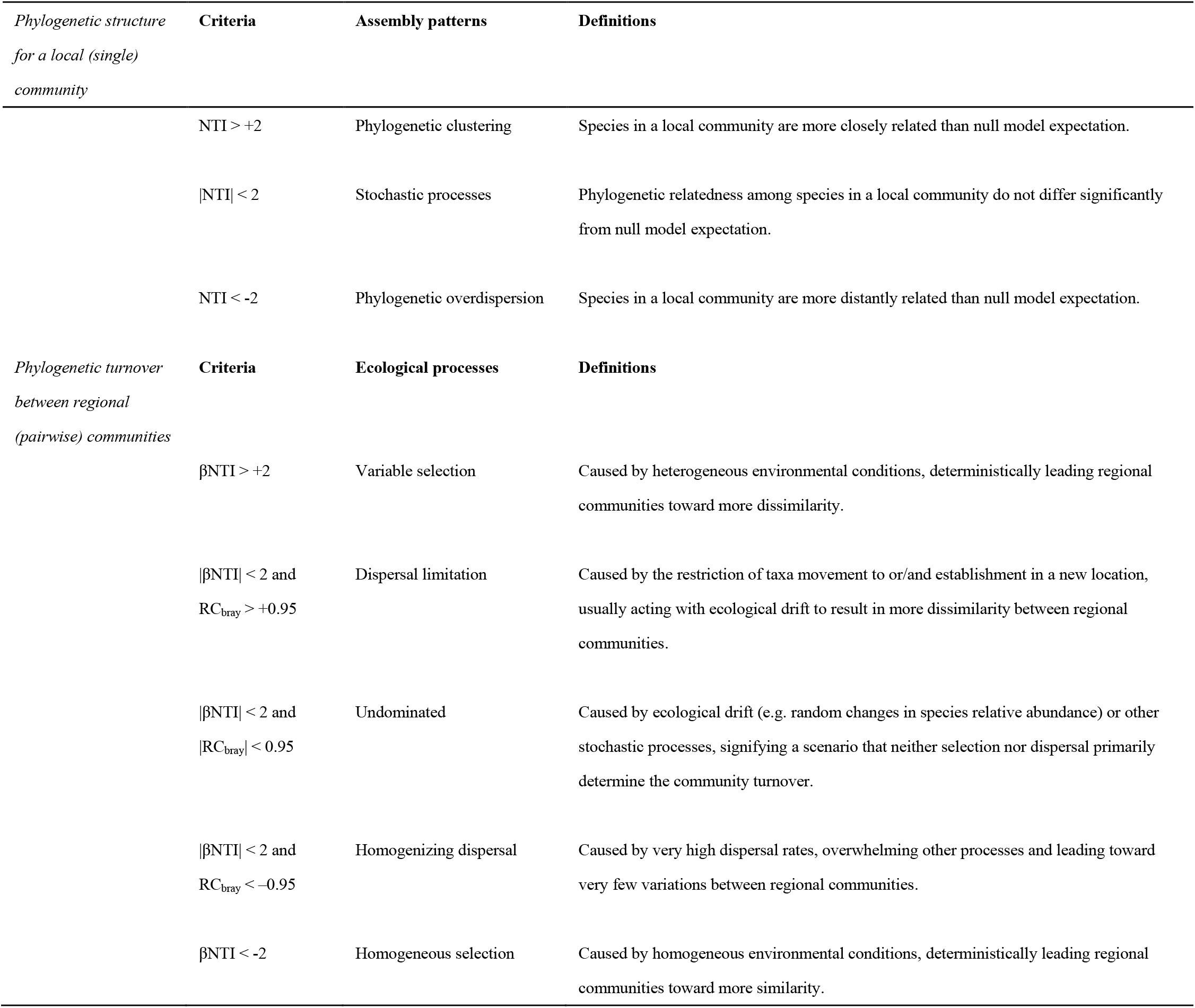
Criteria and definitions for patterns and ecological processes underlying community phylogenetic structures and turnover, respectively (Webb et al., 2002; Stegen et al., 2012; Stegen et al., 2013; Stegen et al., 2015).

Based on these above analyses, we found large differences in vegetation cover, soil properties and microbial community turnover between the early successional stages (ES, including Site I and II) and the later successional stages (LS, including Site III and IV) (see the Result section). In this context, co-occurrence network analysis was performed separately on the data from ES and LS, to reveal microbial co-occurrence patterns along the succession. Two network groups were constructed based on bacterial and fungal communities, respectively. OTUs with occurrence in more than three samples and average relative abundance > 0.1% were selected in early (158 OTUs) and later (168 OTUs) successional stages for bacteria, and in early (83 OTUs) and later (78 OTUs) successional stages for fungi. Spearman correlations between two taxa with adjusted *P*-value (Benjamini and Hochberg, 1995) < 0.01 and rho coefficient > 0.7 or < − 0.7 were considered statistically robust to indicate positive and negative co-occurrences, respectively (Barberan et al., 2012). FDR correction (Benjamini and Hochberg, 1995) of *P*-values was conducted in R using the package ‘vegan’. The taxa involved in the network were designated as co-occurring taxa. A total of 999 Erdős-Rényi random networks (Erdős and Rényi, 1960) with the same number of nodes and edges as each correspondingly observed network were generated using the function *erdos*.*renyi*.*game* in the R package ‘igraph’ (v. 1.2.6). The topological properties and visualizations of networks were performed using Gephi (https://gephi.github.io/). To evaluate the significance of differences in comparisons, Wilcoxon rank sum test and permutational multivariate analysis of variance (PERMANOVA) were conducted in R using the packages ‘stats’ and ‘vegan’. The regression analysis with the method of “loess” (Cleveland and Grosse, 1991) was conducted in R using the packages “ggplot2” (v. 3.3.2).

## 3. Results

### 3.1. Changes in environmental variables along the chronosequence

Vegetation cover and soil properties significantly changed across the post-mining soil chronosequence, especially between the early successional stages (ES, including Site I and II) and the later successional stages (LS, including Site III and IV), but no significant differences were found within ES (Site I *VS* Site II) or LS (Site III *VS* Site IV) (Additional file 1: Table S2 and S3). In ES, the increase of vegetation cover with soil age was accompanied by the significant increase in biomass of bacteria and fungi (Additional file 1: Table S4). In LS, the sites were gradually covered by trees. The soil organic carbon and total nitrogen were significantly higher (ca. two-fold change, *p* < 0.01, by Wilcoxon rank sum test) in LS than those in ES, which was likely attributed to the greater vegetation cover at these mature soil stages (Additional file 1: Table S2). Soil pH showed an opposite pattern, gradually decreasing with soil age from 7.4 to 6.7.

### 3.2. Phylogenetic structures and turnover of microbial communities across the chronosequence

All NTI values of bacterial communities across the chronosequence were greater than + 2 (Fig. 2), indicating that bacterial community structures were more phylogenetically clustered than null model expectation (Stegen et al., 2012). Bacterial NTI values increased with soil age (*p* < 0.05) in ES, thereafter showing less variability in LS (Fig. 2). Bacterial NTI values were significantly positively correlated with vegetation cover but not with soil properties (Additional file 1: Table S5 and S6). Interestingly, bacterial NTI values showed significant positive correlations with bacterial biomass (based on both qPCR and PLFA results) only in ES (Additional file 1: Table S6). NTI values of most fungal communities were greater than + 2 (Fig. 2), indicating that fungal community structures were phylogenetically clustered. Fungal NTI values did not significantly correlate with vegetation cover or fungal biomass (*p* > 0.05), but were only significantly positively correlated with soil pH in LS (Additional file 1: Table S5 and S6).

**Fig. 2.**
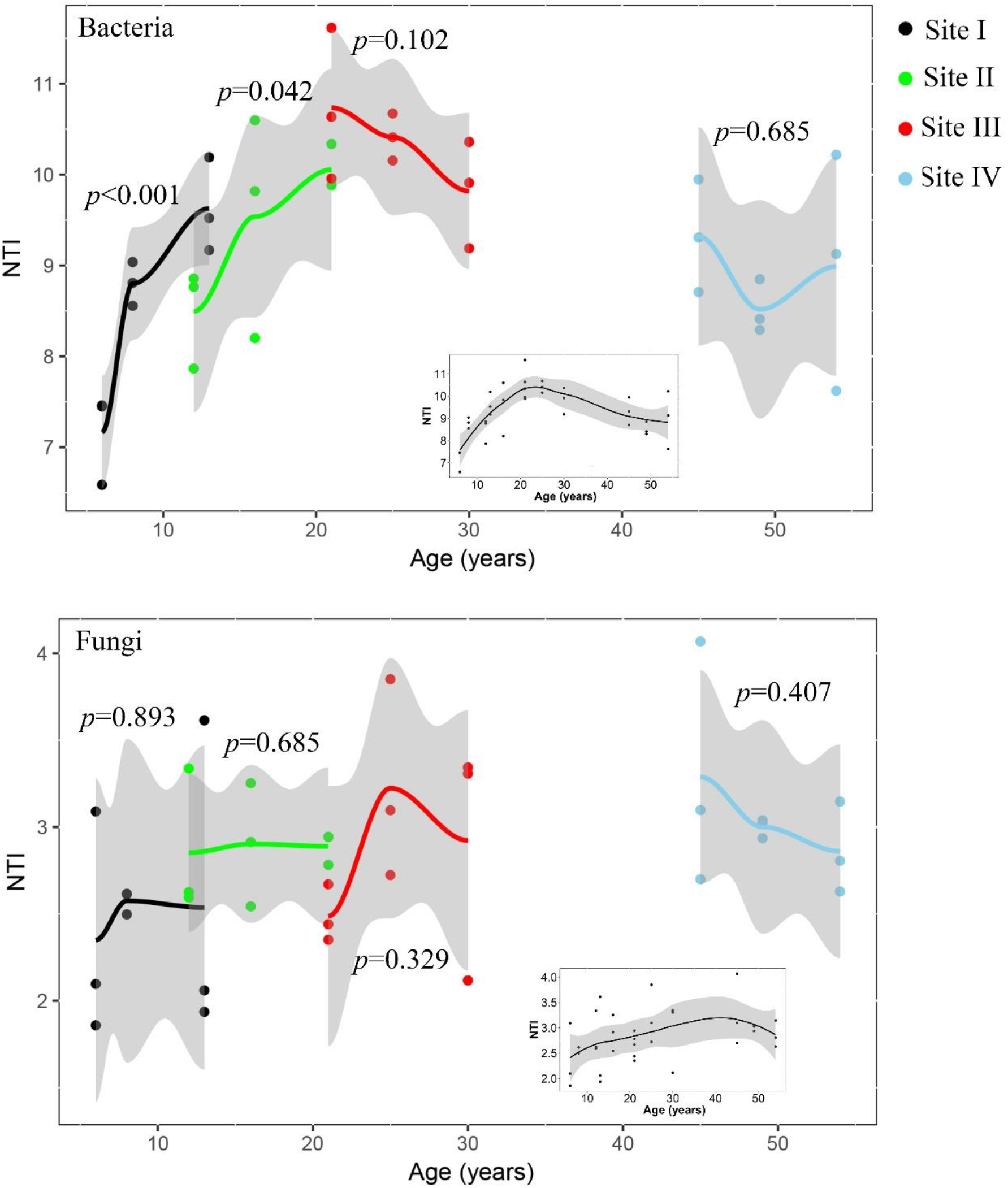
Distribution of NTI values at different sites across the chronosequence, and all sites (embedded). *P*-values of spearman’s correlations between ages and NTI values at different sites are shown. The method of the regression is “loess”. Grey shadow represents 95% confidence intervals.

Across short phylogenetic distances for both bacterial and fungal communities, significant positive correlations were found between OTU phylogenetic distances and OTU niche distances (Additional file 1: Fig. S2), indicating a significant phylogenetic signal. Thus, bacterial and fungal communities exhibited phylogenetic niche conservatism, supporting the use of phylogenetic information to derive ecological inferences (Losos, 2008; Stegen et al., 2013). With the combination of RC_bray_ and βNTI, we quantified the ecological processes governing the turnover of microbial community structures (Fig. 3), based on the criteria shown in Table 1. For the turnover of bacterial community structures, variable selection was stronger in ES (63.9 % at Site I and 69.4 % at Site II) than in LS (2.8 % at Site III and 38.9 % at Site IV), whereas homogeneous selection was stronger in LS (8.3 % at Site III and 30.6 % at Site IV) than in ES (0 % at Site I and 5.6 % at Site II). Across all successional stages (including the turnover between sites and within sites), the turnover of bacterial community structures was mainly driven by dispersal limitation and environmental selection (variable and homogeneous selection) (Fig. 3a). For the turnover of fungal community structures, variable selection was more important in LS (44.4 % at Site III and 52.8 % at Site IV) than in ES (25 % at Site I and 2.7 % at Site II). Across all successional stages, the turnover of fungal community structures was largely driven by dispersal limitation, undominated processes and environmental selection (Fig. 3b).

**Fig. 3.**
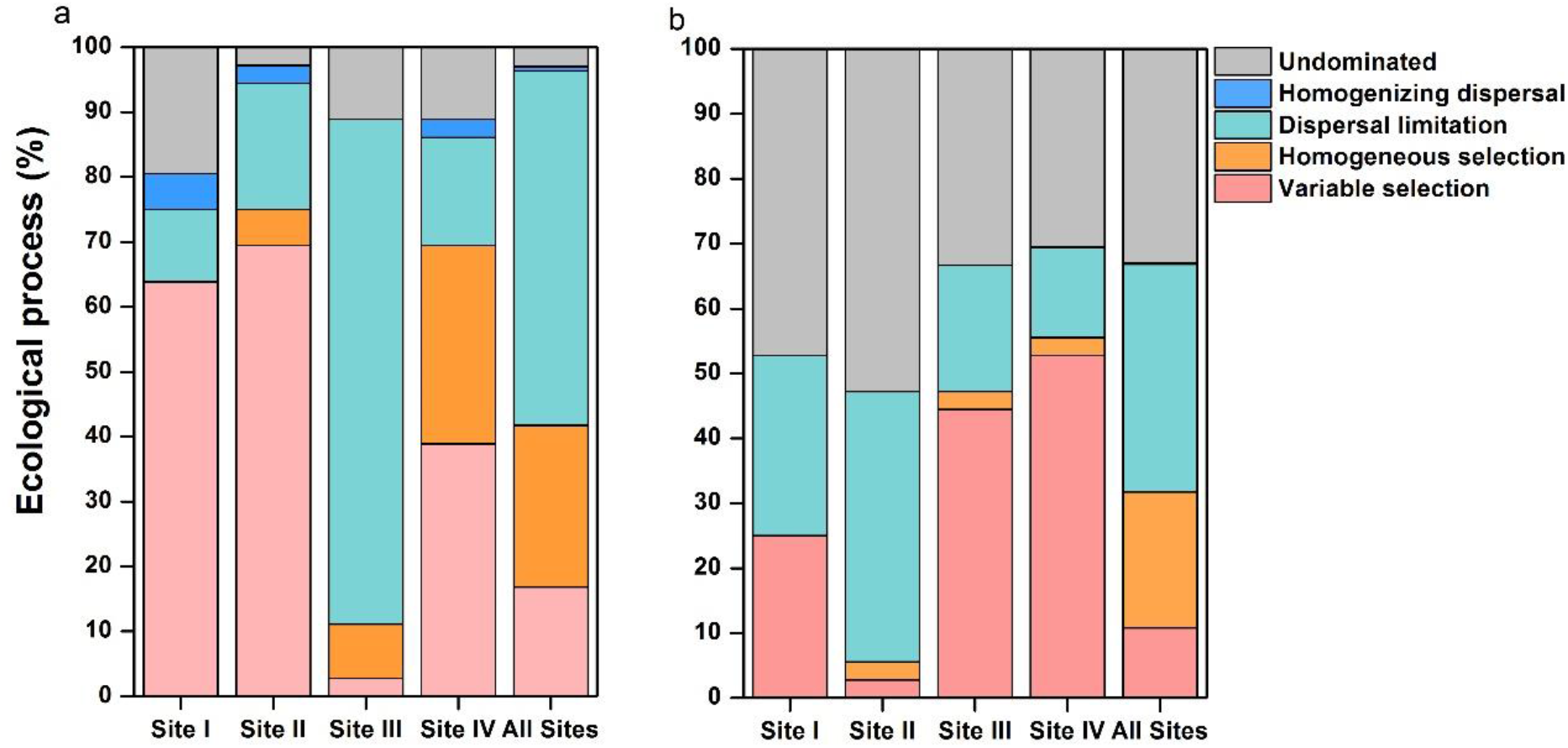
Relative contributions of different ecological processes to bacterial (a) and fungal (b) community turnover at different sites across the soil chronosequence. All Sites: including the community turnover between sites and within sites.

### 3.3. Microbial co-occurrence patterns along succession

Owing to the large differences in vegetation cover, soil properties and microbial community turnover between ES and LS, we constructed networks separately based on data in ES and LS, to reveal the differences in microbial co-occurrence patterns across successional stages. Among the network topological properties, the observed “modularity” was greater than that in the corresponding Erdős-Rényi random networks (Additional file 1: Table S7), suggesting that observed networks had modular structures (Newman, 2006). Other topological properties such as ‘network diameter’ and ‘average path length’ in each observed network were different from those in the corresponding Erdős-Rényi random networks (Additional file 1: Table S7). These implied that all the observed networks were distinguishable from random networks.

All four networks were dominated by co-presence (positive connections) (Fig. 4 and Additional file 1: Table S7). In the bacterial networks, the nodes (OTUs) were mainly from the phyla Proteobacteria, Actinobacteria, Gemmatimonadetes and Bacteroidetes, and node number declined from ES to LS (Fig. 4). The node degree (number of connections to a node) that represents the connectedness of the network significantly declined from ES to LS (Fig. 5a). The network average degree (representing network complexity (Deng et al., 2012; Dini-Andreote et al., 2014)) and the number of total connections also declined from ES to LS (Additional file 1: Fig. S3 and Table S7). Taken together, the signals of network average degree, node degree, node number, and connection number indicated a decrease in the network complexity from ES to LS. Notably, the decrease in the complexity was more obviously reflected in co-presence than in co-exclusion (negative connections) (Fig. 5a and Additional file 1: Fig. S3 and Table S7). However, in the fungal networks, an opposite pattern was observed, such that the network complexity increased from ES to LS (Fig. 4 and 5a, and Additional file 1: Fig. S3 and Table S7).

**Fig. 4.**
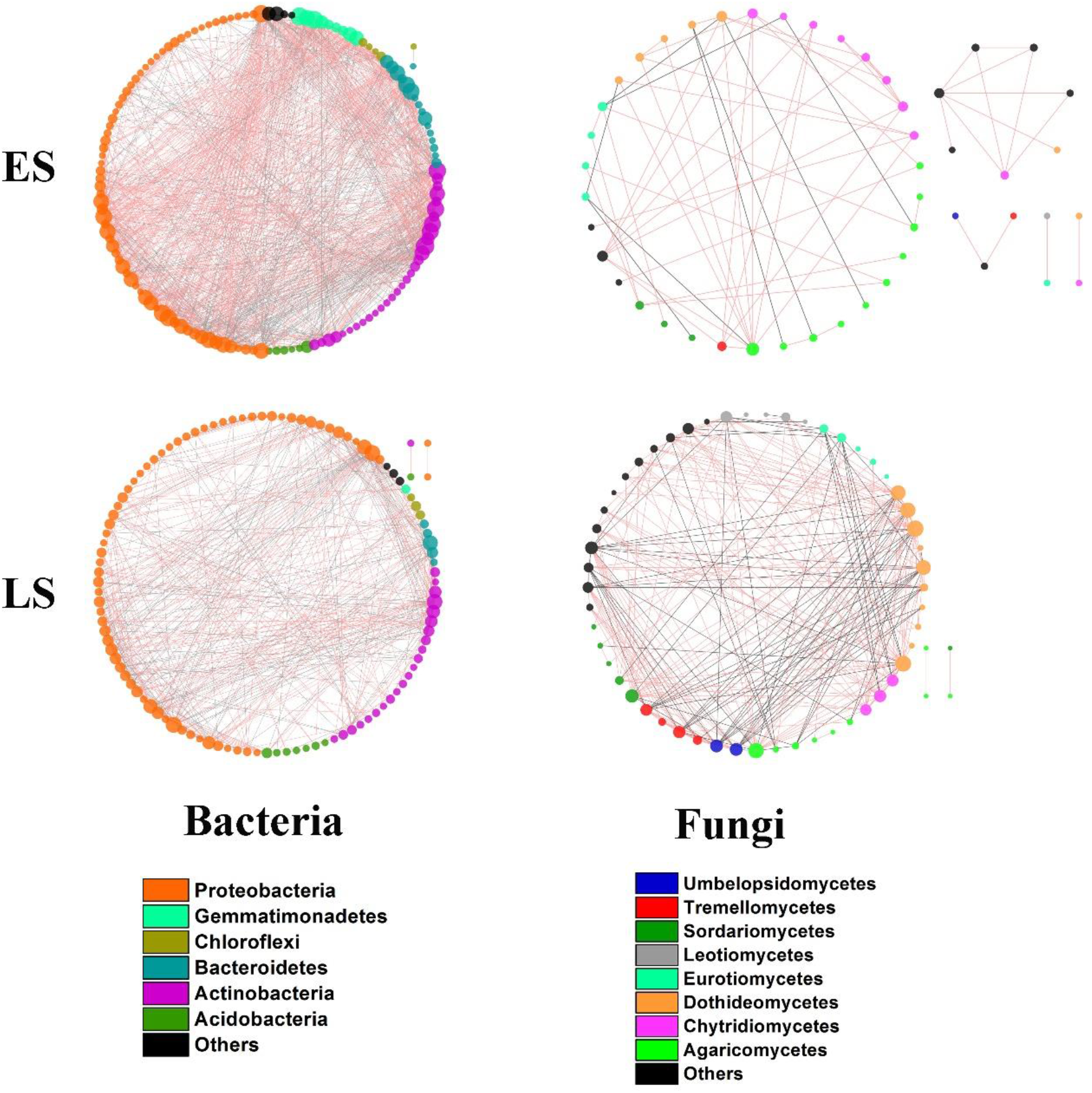
Co-occurrence networks of bacterial and fungal taxa, respectively. The nodes are sized by node degree (number of connections to a node), and colored by bacterial phyla and fungal classes, respectively. Positive and negative connections are colored by red and grey, respectively. ES: early successional stages including Site I and II; LS: later successional stages including Site III and IV.

**Fig. 5.**
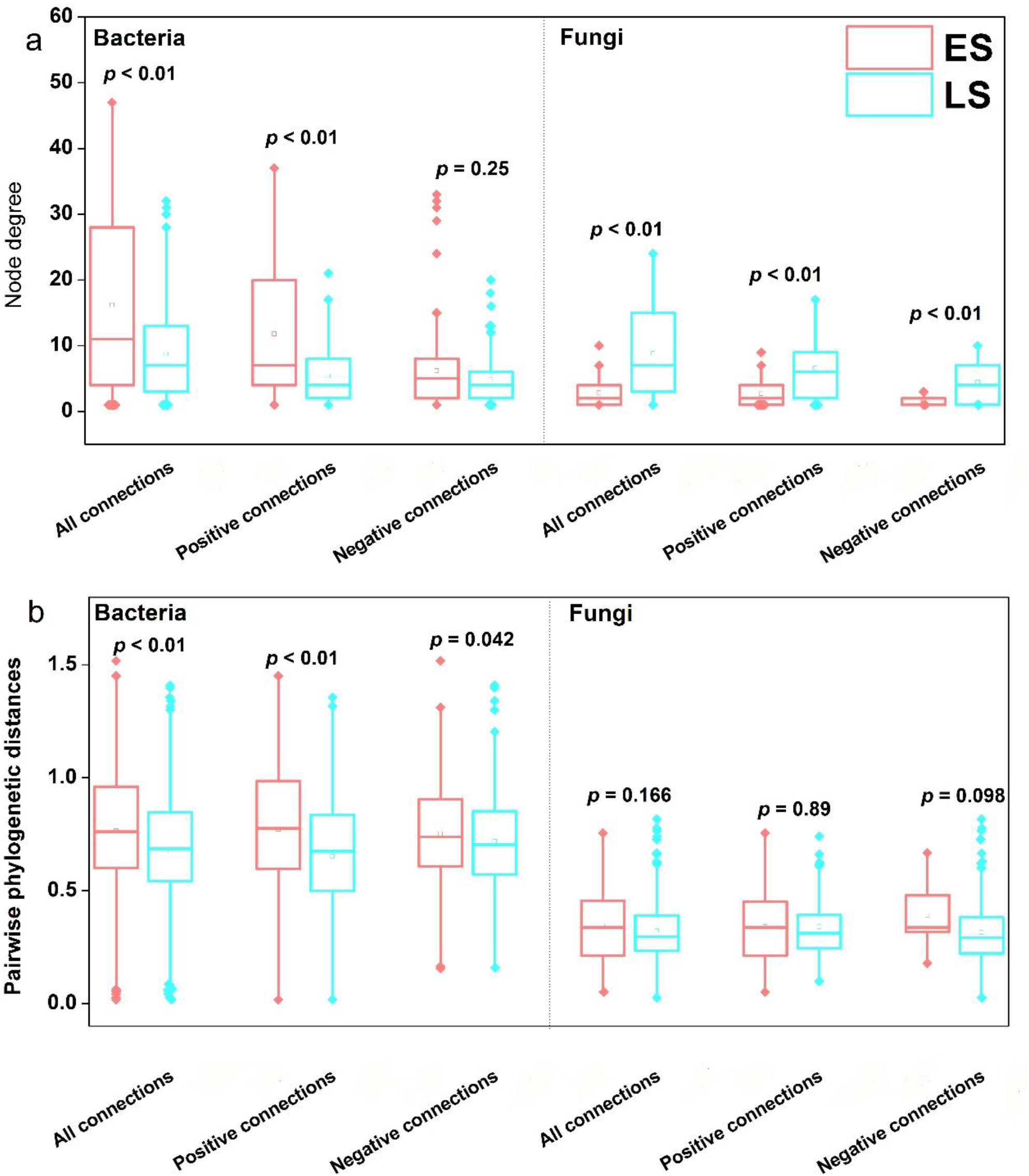
Node degree (a) and pairwise phylogenetic distances between connecting nodes (b) in bacterial and fungal networks. All connections: the network consisting of all connections (edges); positive connections: the sub-network consisting of only positive connections; negative connections: the sub-network consisting of only negative connections. The *P*-value is shown for the statistical significance in each pairwise comparison based on Wilcoxon rank sum test. The square and line inside the box represent the mean and median, respectively. ES: early successional stages including Site I and II; LS: later successional stages including Site III and IV.

Microbial co-occurrence is influenced by microbial niches and thus is usually tightly related to phylogenetic relatedness between co-occurring taxa (Fig. 1b) (Losos, 2008; Stegen et al., 2012). Therefore, the relationship between microbial co-occurrence patterns and corresponding pairwise phylogenetic distances (between co-occurring taxa) was evaluated (Fig. 5b). Greater pairwise phylogenetic distances between two taxa correspond to greater phylogenetical difference between taxa. In bacterial networks, the average pairwise phylogenetic distances were greater (*p* < 0.01) in ES than in LS (Fig. 5b). The differences in average pairwise phylogenetic distances between ES and LS were greater and had higher significance in bacterial co-presence than co-exclusion. For fungal communities, the average pairwise phylogenetic distances were not significantly different (*p* > 0.05) between in ES and in LS, no matter in co-presence or co-exclusion (Fig. 5b). Collectively, the decreasing network complexity corresponded to the decreasing average pairwise phylogenetic distances in bacterial communities across the chronosequence, while there was no such corresponding relationship observed in fungal communities.

There were great variations in microbial co-occurrence patterns across the chronosequence, which was probably related to the turnover of these co-occurring microorganisms (that were involved in networks). Thus, we further evaluated different ecological processes determining the turnover of these co-occurring microorganisms. We found that dispersal limitation played a dominant role in the turnover of co-occurring bacterial communities, and dispersal limitation together with undominated process mainly governed the turnover of co-occurring fungal communities (Additional file 1: Fig. S4). Variable selection exerted a greater role in the turnover of co-occurring bacterial communities in ES than in LS, whereas an opposite pattern was observed in fungi.

## 4. Discussion

### 4.1. Ecological processes driving microbial community succession

The extent of phylogenetic clustering of bacterial communities significantly increased with soil age in ES (Fig. 2), indicating the increasing convergence of bacterial taxa niches. Shifts in microbial community phylogenetic structures are usually related to environmental variations (Stegen et al., 2012; Brown and Jumpponen, 2014). Changes in environmental properties in ES (Additional file 1: Table S2) yielded high environmental heterogeneity, which was further validated by major contributions of variable selection to the turnover of bacterial community structures in ES (Fig. 3a), because variable selection could indicate environmental heterogeneity (Dini-Andreote et al., 2015; Stegen et al., 2015). The high environmental heterogeneity in ES was probably the cause of great dynamics of bacterial community phylogenetic structures. Additionally, significantly positive correlations between bacterial NTI values with vegetation cover in ES (Additional file 1: Table S5) further suggest that environmental selection determined phylogenetic structures of bacterial communities in ES.

Low soil nutrient availability has been reported to enhance phylogenetic clustering of bacterial communities (Feng et al., 2017), which is not consistent with the finding in this study. In ES, the gradually increasing vegetation cover from barren soil via grassland to forest (Additional file 1: Table S2) likely account for the increasing amounts of rhizodeposits and litter, thereby enhancing soil nutrient availability (Harantová et al., 2017). This notion is supported by the increased biomass of bacteria and fungi in ES. Remarkably, the substantial increase of bacterial biomass was accompanied by the increase of bacterial community phylogenetic clustering in ES (Additional file 1: Table S6), which can be explained by taxa heterogeneous responses in our model (Fig. 1a). Specifically, the initial bacterial communities in ES were less phylogenetically clustered (Fig. 2), so the nutrient acquisition strategies among bacterial taxa were presumably less similar (Webb et al., 2002). With progressive increments in nutrient availability along the chronosequence, bacterial taxa performed heterogeneous responses that they substantially expanded their populations over time, with different nutrient acquisition strategies. This notion is supported by the fact that relative abundances of different bacterial genera increased at different rates with soil age in ES (Harantová et al., 2017). Therefore, the substantial but uneven increase of biomass across bacterial taxa probably resulted in subsequent predominance of some taxa and thereby increased phylogenetic clustering of bacterial communities in ES. In prior studies, the increasing phylogenetic clustering of microbial communities was also found to align with the increasing predominance of some taxa in the community subjected to environmental selection (Yan et al., 2016; Wang et al., 2017). This can be explained by that predominant taxa under the similar environmental selection are likely closely-related (Webb et al., 2002), thus resulting in a phylogenetically clustered community. This explanation is supported by evidences that the predominance of well-adapted taxa to pH selection made the community phylogenetically clustered (Tripathi et al., 2018). These examples further prove our inference that less phylogenetically clustered communities caused taxa heterogeneous responses to the increment in nutrient availability and thus co-determined the increasing phylogenetic clustering of bacterial communities with soil age in ES, thereby supporting our conceptual model (Fig. 1a). As succession proceeded, environmental stability and the buffering capacity of soil progressively increased (Dini-Andreote et al., 2014), which probably contributed to the relative stability (less variability) of bacterial community phylogenetic structures in LS. Meanwhile, the significant correlations between bacterial NTI values and vegetation cover in LS indicated that high forest cover probably resulted in homogeneous selection and directly or indirectly contributed to the relative stability of bacterial community phylogenetic structures. Additionally, bacterial homogeneous responses indicated by highly phylogenetic clustering of the bacterial community probably (at least partially) contributed to the relative stability of bacterial community phylogenetic structures in LS.

In ES, the extent of phylogenetic clustering of fungal communities remained relatively stable (less variable) with soil age, despite the increase in fungal biomass. This is consistent with few changes in the relative abundances of most fungal genera across soil ages in ES (Harantová et al., 2017), thus resulting in the relatively low level of community phylogenetic turnover. Besides, the dynamics of fungal community phylogenetic structures in ES were also attributed to the dominance of stochastic processes (e.g., undominated process and dispersal limitation) in community phylogenetic turnover (Fig. 3b). Additionally, neither vegetation cover nor soil properties significantly correlated with fungal NTI values in ES, which further endorses the dominance of stochastic processes in the turnover of fungal community phylogenetic structures.

The relative importance of variable selection in determining fungal community dynamics increased from ES to LS (Fig. 3b), which is not in agreement with other studies where deterministic processes progressively decreased (Tian et al., 2017) or remained unchanged (Brown and Jumpponen, 2015) along successional gradients on glacier forefields. This disagreement was likely due to distinct initial conditions and environmental backgrounds of succession. Soil pH significantly correlated with fungal NTI values in LS. Therefore, the stronger variable selection on fungal community dynamics in LS was probably attributed to soil pH as well as the increased vegetation cover, which is known to at least partially influence fungal communities (Urbanova et al., 2015). Despite the stronger variable selection in LS, fungal community phylogenetic structures did not significantly change with soil age (Fig. 2). This might be explained by important roles of undominated process and dispersal limitation in determining fungal community dynamics (Fig. 3b). Interestingly, in comparison to the ecological processes that determined bacterial community dynamics, we found higher proportions of undominated process driving fungal community dynamics within each site as well as across all sites (Fig. 3). This difference probably explains the distinct dynamics of bacterial and fungal community phylogenetic structures along succession. Dispersal limitation refers to the restriction of taxa movement to and/or establishment in a new location (Martiny et al., 2011; Hanson et al., 2012). In this study, the sampling within a site was conducted in the same geocoordinate with different years (Additional file 1: Table S1). In this context, we assumed that the effect of geographic distance within a site can be neglected, so dispersal limitation (in this case) within each site (Fig. 3) may rather be attributed to the restriction of the continuous establishment of taxa across the chronosequence. Although here we categorized dispersal limitation into stochastic processes, dispersal limitation could be partly caused by deterministic factors (e.g., habitat features that probably changed across the chronosequence in this study) (Hanson et al., 2012). Therefore, the relatively higher proportional influence of dispersal limitation on the dynamics of the bacterial versus fungal communities across all sites (Fig. 3) suggested that bacterial communities were more responsive to the ecological succession than fungal communities.

### 4.2. Mechanisms underlying microbial co-occurrence patterns along succession

Bacterial co-occurrence exhibited decreasing complexity along succession, while the complexity of fungal co-occurrence showed the opposite pattern (Fig. 4 and 5a, and Additional file 1: Fig. S3). A similar development of bacterial co-occurrence was reported in a salt marsh chronosequence, where the higher co-occurrence complexity at initial successional stages was attributed to the higher temporal turnover of bacterial communities (Dini-Andreote et al., 2014). Analogously, in this study, high variability of bacterial community phylogenetic structures (Fig. 2) and high community temporal turnover (Additional file 1: Fig. S5) probably underlay the high complexity of bacterial co-occurrence in ES. Moreover, the importance of variable selection in the turnover of bacterial co-occurring communities and whole communities (Fig. 3 and Additional file 1: Fig. S4) decreased together with bacterial co-occurrence complexity along succession. Because variable selection probably indicates environmental heterogeneity (Dini-Andreote et al., 2015; Stegen et al., 2015), we speculate that the greater environmental heterogeneity contributed to the higher complexity of bacterial co-occurrence in ES.

When linking bacterial co-occurrence patterns to their phylogeny, we found that the decreasing complexity of bacterial co-presence from ES to LS aligned with the general increase of phylogenetic clustering of bacterial communities (Fig. 2), and remarkably with the decrease of average pairwise phylogenetic distances between co-present taxa from ES to LS (Fig. 5b). That is, bacterial co-presence complexity decreased as the co-present taxa became more phylogenetically similar along succession. Based on our model (Fig. 1b), in scenario (1) where taxa co-presence bases on the similar environmental preference between the taxa: if the decrease of bacterial co-presence complexity along succession was attributed to the decreased similarity in environmental preference between the co-present bacterial taxa, the corresponding average pairwise phylogenetic distances between co-present taxa were expected to increase. This would occur because similar environmental preference favors phylogenetically similar (closely related) taxa (Webb et al., 2002; Tripathi et al., 2018). In scenario (2) where taxa co-presence bases on potential cooperation among the taxa: if the decrease of bacterial co-presence complexity along succession was attributed to the decreasing potential cooperation between distantly related bacteria, the corresponding average pairwise phylogenetic distances between co-present taxa were expected to decrease. This would occur because cooperation tends to be established based on metabolic dependencies between distantly related species whose niches are less overlapped (Morris et al., 2013; Zelezniak et al., 2015). In light of these concepts, only the scenario 2 was in agreement with our results, thus indicating that bacterial co-presence was likely an indication of potential cooperation across taxa. Interestingly, this inference, in turn, suggests the tight relationship between potential bacterial interactions and their corresponding phylogenetic relatedness in the natural environment. Most microbial interactions, regardless of cooperation or competition, are mostly established by nutrient demand (Hibbing et al., 2010; Morris et al., 2013). Nutrient addition was reported to substantially alter potential microbial interactions (Banerjee et al., 2016b), and high bacterial co-occurrence complexity has been observed in barren soils (Dini-Andreote et al., 2014; Feng et al., 2017). Low nutrient availability has been suggested to push the exchange of metabolites and nutrients between species for survival (Zelezniak et al., 2015; Morriën et al., 2017). Therefore, in this study, low nutrient availability in ES probably strengthened the potential cooperation between functionally distinct bacteria, which resulted in the relatively higher complexity of bacterial co-presence.

Bacterial co-exclusion can be attributed to potential competition or distinct environmental preferences between co-excluding taxa (Fig. 1b; scenarios 3 and 4). If the decreasing complexity of bacterial co-exclusion from ES to LS (Additional file 1: Fig. S3 and Table S7) indicated the decreasing potential in bacterial competition, the corresponding average pairwise phylogenetic distances between co-excluding taxa were expected to increase, because competition is more prone to occur among phylogenetically similar taxa that occupy a similar niche (Webb et al., 2002; Violle et al., 2011; Stegen et al., 2012). However, this scenario conflicted with our observations (Fig. 5b). If the decreasing complexity of bacterial co-exclusion from ES to LS indicated the decreasing differences in environmental preferences between co-excluding taxa, the corresponding average pairwise phylogenetic distances between co-excluding taxa were expected to decrease, which coincided with our observations (Fig. 5b). This inference was further supported by the increase of phylogenetic clustering of bacterial communities from ES to LS (Fig. 2).

In contrast to bacterial community development, high turnover rates of fungal communities (Additional file 1: Fig. S5) did not result in high network complexity in ES. However, the higher environmental heterogeneity inferred by the higher proportional influence of variable selection on fungal community turnover (Fig. 3b), coincided with the higher complexity of fungal co-occurrence in LS than in ES (Fig.4 and 5a and Additional file 1: Fig. S2). This indicates that the higher environmental heterogeneity may contribute to the higher complexity for fungal co-occurrence in LS. The higher environmental heterogeneity of fungal communities might be due to the increased vegetation cover in LS, as discussed in section 4.1. Interestingly, although fungal co-occurrence complexity increased along succession, the corresponding average pairwise phylogenetic distances between taxa in co-occurrence were not significantly different between in ES and in LS (Fig. 5). Thus, we speculate that fungal co-occurrence patterns along succession were unlikely related to their phylogeny. The less difference in average phylogenetic distances of co-occurring fungi between in ES and in LS was likely a result of the highly similar phylogeny of regional species, which was reflected in the flat dynamics of fungal phylogenetic structures along succession.

## 5. Conclusions and implications

This study shows that bacterial community phylogenetic structures were more responsive than fungal community phylogenetic structures to environmental gradients along the soil primary succession, which was attributed to stronger deterministic effects on bacterial than fungal community phylogenetic turnover. Significant changes in bacterial phylogenetic structures only occurred at early successional stages, and aligned with substantial increase of bacterial biomass. This implies that bacterial taxa niches and bacteria-dependent functions in the ecosystem changed mainly at early successional stages, as the expansion of communities. Both bacterial and fungal co-occurrence patterns significantly varied along succession, but only the former aligned with phylogenetic relatedness between co-occurring taxa, thereby implying potential bacterial cooperation based on our conceptual models. Thus, bacterial co-occurrence patterns along soil primary succession were phylogeny-associated. Taken together, our results boost understanding of ecological processes underlying microbial community development along soil primary succession. Our conceptual models help to address two key fundamental challenges in microbial community assembly and have broad application, as the two challenges exist not only in soil ecosystems but also in other various ecosystems.

## Acknowledgements

We thank Sokolovská Uhelná mining company for providing research permits and background data about the sites.

## Author’s contributions

QL analyzed the data and wrote the manuscript. QL, FDA, TBM, RA, PH, LJL and LM revised the manuscript. LM performed experimental works. JF and PB conceived the study and revised the manuscript. All authors read the manuscript and approved the final draft.

## Data Accessibility

The original sequencing data are available at public database (http://metagenomics.anl.gov/) with dataset number 4741652.3 for bacteria and 4827823.3 for fungi.

## Ethics approval and consent to participate

Not applicable

## Appendices

Supplementary information is available at ##.

