## Additional file 1 for "Microbial phylogenetic relatedness links to distinct successional patterns of bacterial and fungal communities"

**Running title:** Ecological succession of bacteria and fungi

[a] Biology Centre of the Czech Academy of Sciences, Institute of Soil Biology & SoWa Research Infrastructure, Na Sádkách 7, CZ, 37005, České Budějovice, Czech Republic

[b] Department of Plant Science, The Pennsylvania State University, University Park, PA, USA

[c] Huck Institutes of the Life Sciences, The Pennsylvania State University, University Park, PA, USA

[d] Laboratory of Environmental Microbiology, Institute of Microbiology of the Czech Academy of Sciences, Vídeňská 1083, 14220 Praha 4, Czech Republic

[e] Institute for environmental studies, Faculty of Science, Charles University, Benátská 2, 12800, Praha 2, Czech Republic

[f] Department of Geosciences and Natural Resource Management, Faculty of Science, University of Copenhagen, Rolighedsvej 23, 1958 Frederiksberg C, Denmark

[g] Engineering Research Center of Soil Remediation of Fujian Province University, College of Resources and Environment, Fujian Agriculture and Forestry University, Fuzhou 350002,

China

**Table S1.** Information of field sites.

| | Successional age (Sampling time)* | Location | Geocoordinates |
|---|---|---|---|
| Site I | 6 years (2008), 8 years (2010) and 13 years (2015) | Velká podkrušnohorská spoil heap in the Sokolov lignite mining district of Czech Republic | 50.2389711N, 12.6604986E |
| Site II | 12 years (2006), 16 years (2010) and 21 years (2015) | | 50.2319303N, 12.6401353E |
| Site III | 21 years (2006), 25 years (2010) and 30 years (2015) | | 50.2420728N, 12.6793169E |
| Site IV | 45 years (2006), 49 years (2010) and 54 years (2015) | | 50.2551353N, 12.7354503E |

*All samplings are conducted in the end of May.

**Table S2.** Vegetation cover and soil properties across the chronosequence.

| Age(years)/site | Mosses cover (%) | Herbs cover (%) | Shrubs cover (%) | Trees cover (%) | Soil pH | Soil total nitrogen (%) | Soil organic carbon (%) |
|---|---|---|---|---|---|---|---|
| 6/I | 0.0 ± 0.0 a | 9.67 ± 1.86 cd | 0.0 ± 0.0 a | 0.0 ± 0.0 c | 7.40 ± 0.05 | 0.28 ± 0.05 | 4.83 ± 0.69 |
| 8/I | 1.33 ± 0.88 a | 32.67 ± 1.76 bc | 0.0 ± 0.0 a | 0.0 ± 0.0 c | 7.54 ± 0.04 | 0.17 ± 0.01 | 3.57 ± 0.15 |
| 12/II | 1.67 ± 1.67 a | 31.67 ± 3.33 bc | 0.0 ± 0.0 a | 0.0 ± 0.0 c | 6.93 ± 0.04 | 0.17 ± 0.01 | 3.20 ± 0.64 |
| 13/I | 2.00 ± 1.53 a | 25.00 ± 2.89 bcd | 0.0 ± 0.0 a | 0.0 ± 0.0 c | 7.54 ± 0.11 | 0.22 ± 0.01 | 3.87 ± 0.18 |
| 16/II | 9.00 ± 3.00 a | 23.33 ± 4.18 bcd | 1.33 ± 1.33 a | 0.0 ± 0.0 c | 7.54 ± 0.17 | 0.22 ± 0.01 | 2.99 ± 0.47 |
| 21/II | 18.33 ± 7.26 a | 23.33 ± 8.99 bcd | 5.00 ± 5.00 a | 0.0 ± 0.0 c | 6.83 ± 0.23 | 0.28 ± 0.03 | 4.65 ± 0.44 |
| 21/III | 25.00 ± 5.00 a | 3.33 ± 0.17 d | 0.0 ± 0.0 a | 36.67 ± 10.14 b | 6.34 ± 0.08 | 0.33 ± 0.02 | 10.00 ± 1.73 |
| 25/III | 10.67 ± 2.96 a | 2.67 ± 0.73 d | 0.0 ± 0.0 a | 73.33 ± 3.33 a | 6.82 ± 0.29 | 0.53 ± 0.16 | 8.47 ± 1.37 |
| 30/III | 6.00 ± 0.00 a | 3.00 ± 1.00 d | 0.0 ± 0.0 a | 73.33 ± 4.41 a | 6.65 ± 0.39 | 0.49 ± 0.06 | 7.37 ± 1.38 |
| 45/IV | 10.60 ± 9.70 a | 43.33 ± 3.33 ab | 0.0 ± 0.0 a | 3.33 ± 1.67 c | 6.22 ± 0.02 | 0.46 ± 0.07 | 7.57 ± 0.58 |
| 49/IV | 12.00 ± 11.50 a | 28.33 ± 6.01 bc | 0.0 ± 0.0 a | 10.00 ± 5.77 c | 6.25 ± 0.18 | 0.38 ± 0.01 | 6.96 ± 0.24 |
| 54/IV | 18.67 ± 15.71 a | 60.00 ± 8.66 a | 1.00 ± 1.00 a | 17.33 ± 8.76 bc | 6.65 ± 0.36 | 0.40 ± 0.08 | 6.20 ± 1.14 |

Data are expressed as the mean ± SEM. Different letters in column denote statistically significant differences ($p < 0.05$). These data are based on the previous study (Harantová et al., 2017).

**Table S3.** Statistical differences of vegetation cover and soil properties among different sites assessed by PERMANOVA.

| | Vegetation | | Soil properties | |
|---|---|---|---|---|
| | $R^2$ | *P* | $R^2$ | *P* |
| Site I *VS* Site II | 0.15062 | 0.0508 | 0.10188 | 0.1844 |
| Site I *VS* Site III | 0.76686 | 3.00E-04 | 0.67105 | 1.00E-04 |
| Site I *VS* Site IV | 0.33971 | 3.00E-04 | 0.70598 | 3.00E-04 |
| Site II *VS* Site III | 0.71449 | 2.00E-04 | 0.63572 | 1.00E-04 |
| Site II *VS* Site IV | 0.21598 | 0.002 | 0.63409 | 2.00E-04 |
| Site III *VS* Site IV | 0.65756 | 3.00E-04 | 0.12221 | 0.1328 |
| ES *VS* LS | 0.33982 | 1.00E-04 | 0.62124 | 1.00E-04 |

ES: early successional stages including Site I and II; LS: later successional stages including Site III and IV.

**Table S4.** Bacterial and fungal biomass based on qPCR and phospholipid fatty acid, respectively, along the succession.

| | Bacterial rDNA ($10^6$ copies $g^{-1}$ soil dry mass) | Fungal rDNA ($10^6$ copies $g^{-1}$ soil dry mass) | Bacterial PLFA (nmol $g^{-1}$) | Fungal PLFA (nmol $g^{-1}$) |
|---|---|---|---|---|
| Site I6 | 12.29 ± 4.38 b | 0.43 ± 0.12 c | 5.25 ± 1.82 e | 0.56 ± 0.19 e |
| Site I8 | 43.99 ± 3.34 b | 2.64 ± 0.22 c | NA. | NA. |
| Site I13 | 234.19 ± 12.64 b | 19.91 ± 1.84 bc | 13.98 ± 4.28 de | 1.63 ± 0.39 de |
| Site II12 | 200.76 ± 12.42 b | 22.33 ± 2.68 bc | 15.68 ± 10.42 cde | 3.55 ± 2.49 cde |
| Site II16 | 143.95 ± 19.81b | 12.10 ± 3.23 c | NA. | NA. |
| Site II21 | 486.62 ± 102.98 b | 58.86 ± 12.83 bc | 28.12 ± 6.66 bcd | 5.95 ± 1.88 bcd |
| Site III21 | 1,935.00 ± 1,017.85 a | 241.49 ± 92.11 a | 82.67 ± 5.33 a | 21.96 ± 1.56 a |
| Site III25 | 407.32 ± 87.93 b | 45.15 ± 10.03 bc | NA. | NA. |
| Site III30 | 935.49 ± 251.42 ab | 114.59 ± 20.31 abc | 52.20 ± 20.61 abc | 8.81 ± 5.24 abc |
| Site IV45 | 1,023.74 ± 120.57 ab | 218.97 ± 24.65 a | 58.27 ± 9.95 ab | 9.88 ± 2.26 ab |
| Site IV49 | 1,198.72 ± 44.18 ab | 172.37 ± 9.94 ab | NA. | NA. |
| Site IV54 | 1,190.31 ± 242.91 ab | 138.16 ± 33.60 abc | 6.46 ± 2.07 ab | 9.06 ± 2.36 a |

Data are expressed as the mean ± SEM. These data are based on the previous study (Harantová et al., 2017). NA: not assessed. Different letters in column denote statistically significant differences ($p < 0.05$).

**Table S5.** Mantel test for Spearman's correlations between Euclidean distances of NTI values and vegetation cover, through 9999 permutations.

| | | R | *P* |
|---|---|---|---|
| ES | Bacterial NTI | 0.3421 | **0.005** |
| LS | Bacterial NTI | 0.1723 | **0.0379** |
| All stages | Bacterial NTI | 0.1461 | **0.0322** |
| ES | Fungal NTI | -0.0727 | 0.701 |
| LS | Fungal NTI | -0.0247 | 0.5545 |
| All stages | Fungal NTI | 0.0125 | 0.41 |

ES: early successional stages including Site I and II; LS: later successional stages including Site III and IV.

**Table S6.** Spearman's correlations between NTI and soil properties and biomass.

| | | pH | Total nitrogen | Organic carbon | Biomass (qPCR) | Biomass (PLFA) |
|---|---|---|---|---|---|---|
| ES | Bacterial NTI | -0.057 | 0.191 | -0.013 | 0.748** | 0.685* |
| LS | Bacterial NTI | 0.449 | -0.36 | 0.086 | -0.583* | -0.056 |
| All stages | Bacterial NTI | -0.12 | 0.2 | 0.275 | 0.211 | 0.440* |
| ES | Fungal NTI | -0.433 | 0.259 | 0.189 | 0.239 | 0.221 |
| LS | Fungal NTI | 0.593** | -0.179 | -0.457 | -0.579* | -0.567 |
| All stages | Fungal NTI | -0.158 | 0.255 | 0.179 | 0.287 | 0.086 |

** Significant at $p < 0.01$, * $p < 0.05$. ES: early successional stages including Site I and II; LS: later successional stages including Site III and IV.

**Table S7.** Network topological properties.

| | | All associations | | | | | Positive associations | | Negative associations | |
|---|---|---|---|---|---|---|---|---|---|---|
| | | Average degree | Network diameter | Graph density | Modularity | Average path length | Edge number | Node number | Edge number | Node number |
| Bacteria | ES | 16.2 | 8 | 0.117 | 0.31 | 2.822 | 772 | 131 | 362 | 117 |
| | Corresponding random network | 16.2 | 3 | 0.117 | 0.197 | 2.016 | NA | NA | NA | NA |
| | LS | 8.793 | 9 | 0.079 | 0.392 | 3.094 | 291 | 108 | 194 | 80 |
| | Corresponding random network | 8.793 | 4 | 0.079 | 0.276 | 2.394 | NA | NA | NA | NA |
| Fungi | ES | 2.833 | 8 | 0.060 | 0.632 | 3.487 | 59 | 44 | 9 | 12 |
| | Corresponding random network | 2.833 | 4 | 0.060 | 0.451 | 1.796 | NA | NA | NA | NA |
| | LS | 8.842 | 6 | 0.158 | 0.342 | 2.437 | 174 | 53 | 78 | 35 |
| | Corresponding random network | 8.842 | 4 | 0.158 | 0.273 | 2.053 | NA | NA | NA | NA |

NA means non-calculable. ES: early successional stages including Site I and II; LS: later successional stages including Site III and IV. Means of 999 Erdős-Rényi random networks are shown.

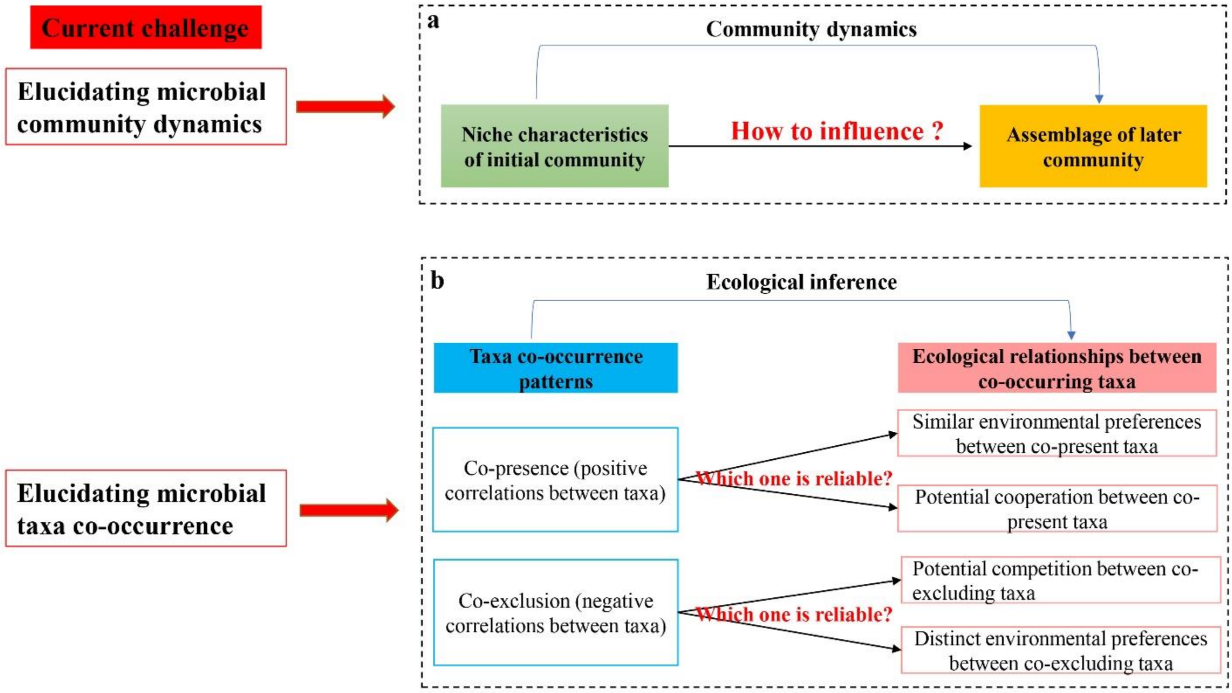


**Fig. S1** Current challenges in elucidating microbial community dynamics (a) and microbial taxa co-occurrence patterns (b), respectively. In **Fig. S1a**, community dynamics is the transition from an initial community to a later community. However, it remains challenging to understand how the niche characteristics of the initial community would affect community dynamics. In **Fig. S1b**, one specific co-occurrence patterns could imply two distinct ecological inferences, so it remains challenging to achieve reliable ecological understanding of microbial co-occurrence. These two key fundamental challenges hinder our understanding of mechanisms underlying microbial community assembly.

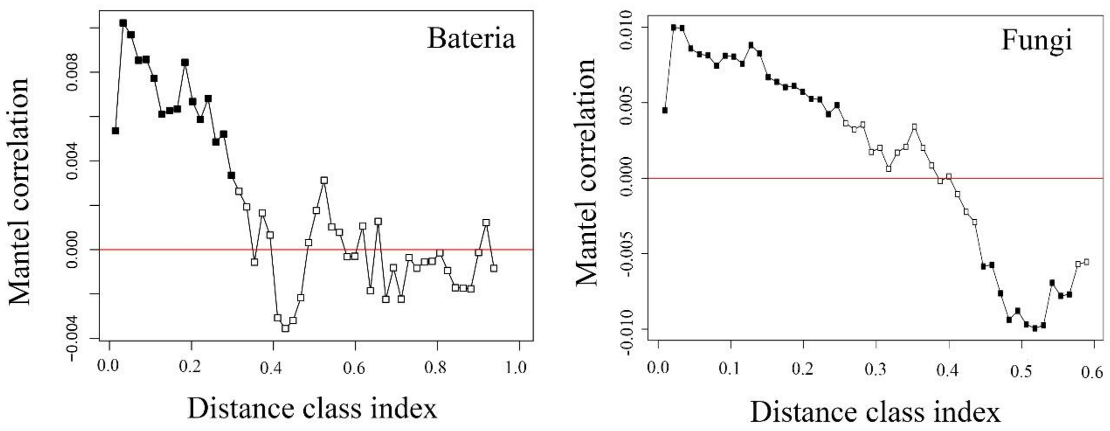


**Fig. S2** Pearson correlation resulting from Mantel correlogram between the pairwise matrix of OTU phylogenetic distances and OTU niche distances with 999 permutations. Solid and open symbols denote significant ($p < 0.05$) and non-significant correlations, respectively.

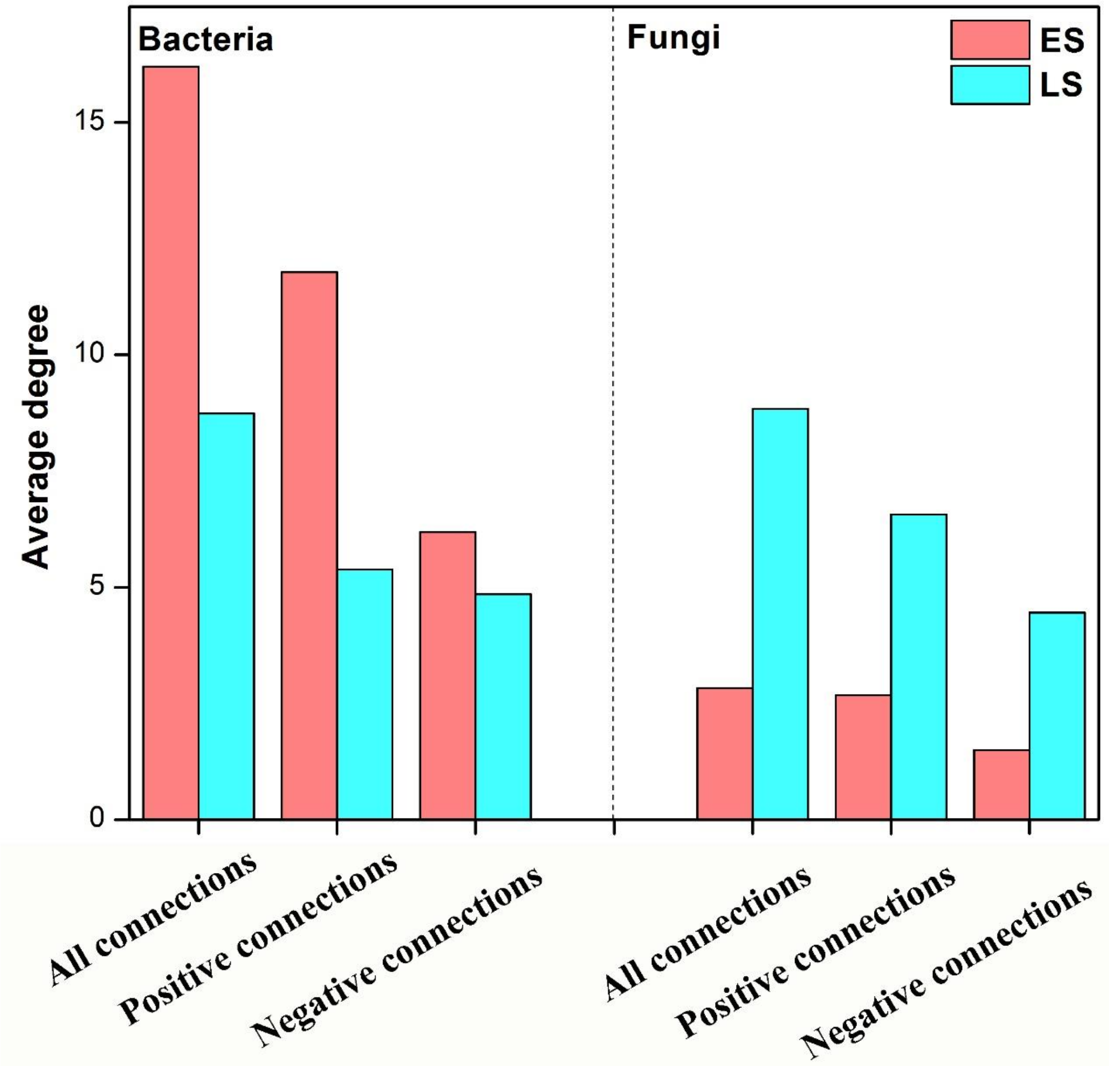


**Fig. S3** The average degree distributions in two network groups, respectively. ES: early successional stages including Site I and II; LS: later successional stages including Site III and IV. All connections: the network consisting of all connections (edges); positive connections: the sub-network consisting of only positive connections; negative connections: the sub-network consisting of only negative connections.

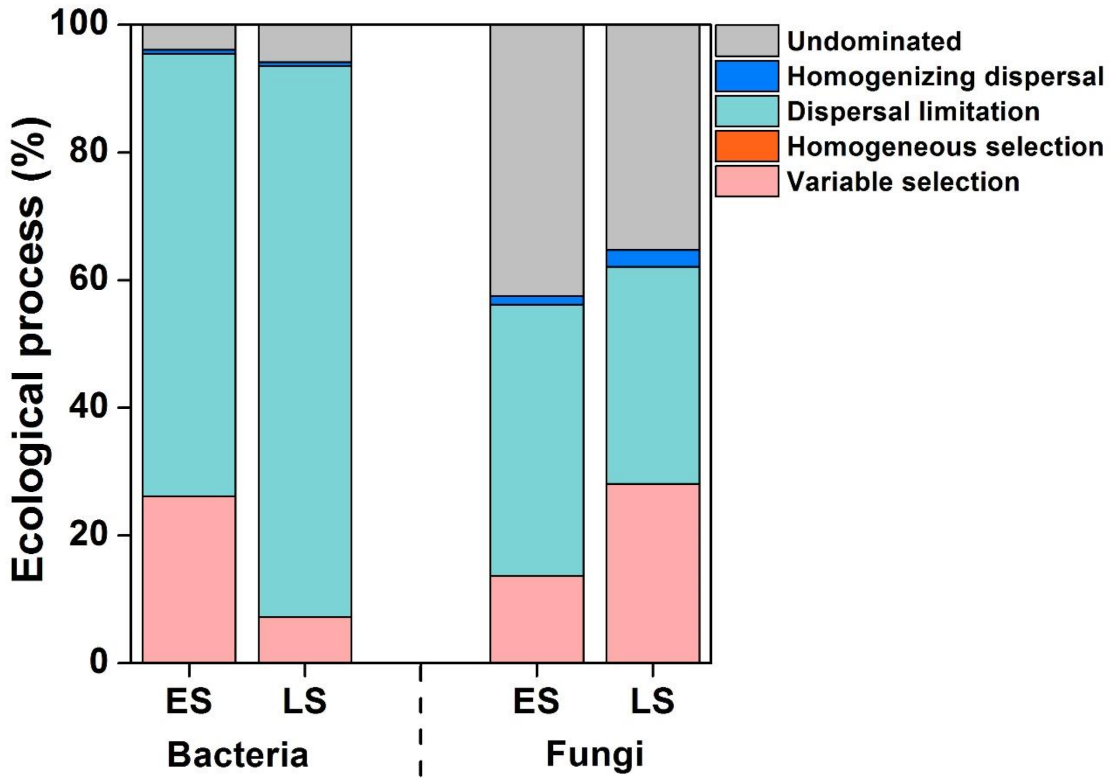


**Fig. S4** Relative contributions of different ecological processes to the turnover of co-occurring bacterial (a) and fungal (b) community, respectively, at different sites across chronosequence. ES: early successional stages including Site I and II; LS: later successional stages including Site III and IV.

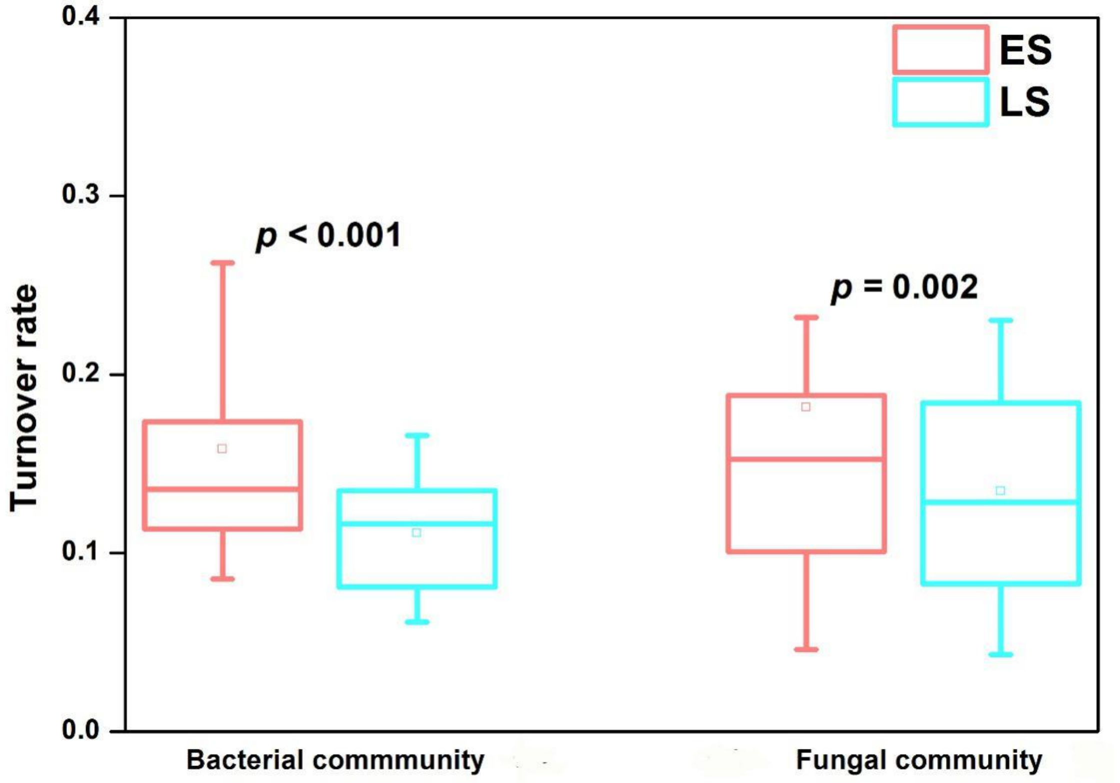


**Fig. S5** Turnover rates of bacterial and fungal communities in ES (including Site I and II) and LS (including Site III and IV). The pairwise Bray-Curtis dissimilarity is divided by the corresponding time interval to evaluate the turnover rates. The *P*-value is shown for the statistical significance in each pairwise comparison based on Wilcoxon rank sum test. The square and line inside the box represent mean and median, respectively.
