## Additional file 2 for "Microbial phylogenetic relatedness links to distinct successional patterns of bacterial and fungal communities"

**Running title:** Ecological succession of bacteria and fungi

[a] Biology Centre of the Czech Academy of Sciences, Institute of Soil Biology & SoWa Research Infrastructure, Na Sádkách 7, CZ, 37005, České Budějovice, Czech Republic

[b] Department of Plant Science, The Pennsylvania State University, University Park, PA, USA

[c] Huck Institutes of the Life Sciences, The Pennsylvania State University, University Park, PA, USA

[d] Laboratory of Environmental Microbiology, Institute of Microbiology of the Czech Academy of Sciences, Vídeňská 1083, 14220 Praha 4, Czech Republic

[e] Institute for environmental studies, Faculty of Science, Charles University, Benátská 2, 12800, Praha 2, Czech Republic

[f] Department of Geosciences and Natural Resource Management, Faculty of Science, University of Copenhagen, Rolighedsvej 23, 1958 Frederiksberg C, Denmark

[g] Engineering Research Center of Soil Remediation of Fujian Province University, College of Resources and Environment, Fujian Agriculture and Forestry University, Fuzhou 350002,

China

**S1.** DNA extraction.

DNA extraction followed a modified method of Miller (Sagova-Mareckova et al., 2008). Soil (180 mg) was pretreated with 1 M $CaCO_3$ solution (1:1 ratio by volume) for 1 hour under room temperature. A buffer (600 µl, 50 mM Na-phosphate buffer [pH 8], 50 mM NaCl, 500 mM Tris-HCl [pH 8], and 5% sodium dodecyl sulfate) and 300 µl of phenol-chloroform–isoamyl alcohol (25:24:1) and 0.5 g sterile glass beads (0.1-mm diameter [0.25 mg] and 0.5-mm diameter [0.25 mg]) were added to the above mixture of soil and $CaCO_3$, and then was homogenized in a beater for 45 s, at 2,500 rpm, twice. The homogenate was centrifuged at 16,000 × g for 2 min. The supernatant was mixed with the same volume of phenol-chloroform–isoamyl alcohol (25:24:1) and centrifuged at 6,000 × g for 5 min. The supernatant was mixed with an equal volume of chloroform-isoamyl alcohol (24:1) and centrifuged at 16,000 × g for 5 min. To the supernatant, NaCl was added to a final concentration of 1.5 M, and CTAB was added to 1% and incubated at 65°C for 30 min. The incubated solution was cooled, mixed with an equal volume of chloroform-isoamyl alcohol (24:1), and centrifuged at 3,400 × g for 20 min. The supernatant was then precipitated with isopropanol. Geneclean Turbo Kit (Biogenic) was used to purify DNA extracts, following the manufacturer's instructions.

**S2.** Phylogenetic null model.

Generally, the calculation of phylogenetic null model followed previous methods (Stegen et al., 2013; Stegen et al., 2015). Briefly, for bacteria and fungi, respectively, we calculated a phylogenetic null model expectation by a phylogenetic randomization procedure within each site (to evaluate the turnover of communities within one site) and across sites (to evaluate the turnover of communities during the whole succession process), respectively. During the randomization procedure, species names and abundances were shuffled across the tips of the phylogeny, then βMNTD was recalculated to provide a null value, and finally a null distribution of βMNTD was generated by repeating the randomization 999 times. The degree to which observed βMNTD deviates from the mean of the null distribution βMNTD normalized by its standard deviation is referred to as the β-nearest taxon index (βNTI).
